# Prediction of cancer treatment response from histopathology images through imputed transcriptomics

**DOI:** 10.1101/2022.06.07.495219

**Authors:** Danh-Tai Hoang, Gal Dinstag, Leandro C. Hermida, Doreen S. Ben-Zvi, Efrat Elis, Katherine Caley, Stephen-John Sammut, Sanju Sinha, Neelam Sinha, Christopher H. Dampier, Chani Stossel, Tejas Patil, Arun Rajan, Wiem Lassoued, Julius Strauss, Shania Bailey, Clint Allen, Jason Redman, Tuvik Beker, Peng Jiang, Talia Golan, Scott Wilkinson, Adam G. Sowalsky, Sharon R. Pine, Carlos Caldas, James L. Gulley, Kenneth Aldape, Ranit Aharonov, Eric A. Stone, Eytan Ruppin

## Abstract

Advances in artificial intelligence have paved the way for leveraging hematoxylin and eosin (H&E)-stained tumor slides for precision oncology. We present ENLIGHT-DeepPT, an approach for predicting response to multiple targeted and immunotherapies from H&E-slides. In difference from existing approaches that aim to predict treatment response directly from the slides, ENLIGHT-DeepPT is an indirect two-step approach consisting of (1) DeepPT, a new deep-learning framework that predicts genome-wide tumor mRNA expression from slides, and (2) ENLIGHT, which predicts response based on the DeepPT inferred expression values. DeepPT successfully predicts transcriptomics in all 16 TCGA cohorts tested and generalizes well to two independent datasets. Importantly, ENLIGHT-DeepPT successfully predicts true responders in five independent patients’ cohorts involving four different treatments spanning six cancer types with an overall odds ratio of 2.44, increasing the baseline response rate by 43.47% among predicted responders, without the need for any treatment data for training. Furthermore, its prediction accuracy on these datasets is comparable to a supervised approach predicting the response directly from the images, trained and tested on the same cohort in cross validation. Its future application could provide clinicians with rapid treatment recommendations to an array of different therapies and importantly, may contribute to advancing precision oncology in developing countries.

**Statement of Significance:** ENLIGHT-DeepPT is the first approach shown to successfully predict response to *multiple* targeted and immune cancer therapies from H&E slides. In distinction from all previous H&E slides prediction approaches, it does not require supervised training on a specific cohort for each drug/indication treatment but is trained to predict expression on the TCGA cohort and then can predict response to an array of treatments without any further training. ENLIGHT-DeepPT can provide rapid treatment recommendations to oncologists and help advance precision oncology in underserved regions and low-income countries.

## INTRODUCTION

Histopathology has long been considered the gold standard of clinical diagnosis and prognosis in cancer. In recent years, the use of tumour molecular profiling within the clinic has allowed for more accurate cancer diagnostics, as well as the delivery of precision oncology [1–3]. Rapid advances in digital histopathology have also allowed the extraction of clinically relevant information embedded in tumor slides by applying machine learning and artificial intelligence methods, capitalizing on recent advancements in image analysis via deep learning [4]. Key advances are already underway, as whole slide images (WSI) of tissue stained with hematoxylin and eosin (H&E) have been used to computationally diagnose tumors [5–8], classify cancer types [7,9–13], distinguish tumors with low or high mutation burden [14], identify genetic mutations [6,15–23], predict patient survival [24–29], detect DNA methylation patterns [30] and mitosis [31], and quantify tumor immune infiltration [32].

Previous work has already impressively unravelled the potential of harnessing next-generation digital pathology to predict response to therapies *directly* from images [33–37]. In these direct supervised learning approaches, predicting response to therapy directly from the WSI requires large datasets consisting of matched imaging and response data. As such, they require a specific cohort for each drug/indication treatment that is to be predicted. However, the availability of such data on a large scale is still fairly limited, restricting the applicability of this approach and raising concerns about the generalizability of supervised predictors to other cohorts.

To overcome this challenge, we turned to develop and study a generic methodology for generating WSI-based predictors of patients’ response for a broad range of cancer types and therapies, *which does not require matched WSI and response datasets for training*. To accomplish this, we have taken an *indirect* two-step approach: First, we developed DeepPT (<u>Deep</u> <u>P</u>athology for <u>T</u>ranscriptomics), a novel deep-learning framework for imputing (predicting) gene expression from H&E slides, which extends upon previous valuable work on this topic [38–43]. The DeepPT models are cancer type-specific and are built by training on matched WSI and expression data from the TCGA. Second, given gene expression values predicted by these models for a new patient, we apply our previously published approach, ENLIGHT [44], originally developed to predict patient response from *measured* tumor transcriptomics, to predict response from the DeepPT *imputed* transcriptomics.

We proceed to provide an overview of DeepPT’s architecture and a brief recap of ENLIGHT’s workings, the study design and the cohorts analysed. We then describe the results obtained, in each of the two steps of ENLIGHT-DeepPT. First, we study the ability to predict tumor expression, showing the performance of the trained DeepPT models in predicting the gene bulk expression in 16 TCGA cohorts and in two independent, unseen cohorts. Second, we analyze five independent clinical trial datasets of patients with different cancer types that were treated with various targeted and immune therapies. Critically, those are test cohorts, on which DeepPT was never trained. We show that ENLIGHT, adhering to the parameters used in its original publication [44] without any adaptation, can successfully predict the true responders from the expression values imputed by DeepPT, using only H&E images. We then compare its prediction accuracy to that of a direct approach that predicts the response directly from the images. Overall, our results show that combining digital pathology with an expression-based response prediction approach offers a promising new way to provide clinicians with almost immediate treatment recommendations that may help guide patients’ treatment until more information arrives from multi-omics biomarker screens.

Finally, and importantly, we believe that *ENLIGHT-DeepPT* may make precision oncology more accessible to patients in low- and middle-income countries (LMICs) and other under-served regions. As have been noted recently, LMICs now have increasing access to cancer medicines, primarily due to the WHO’s expansion of the list of essential medicines (EMLs), and different partnerships [45]. However, these encouraging developments have not yet been matched by access to cancer diagnostics, which are obviously needed to provide precision treatments as beneficial as possible. As noted in [45], there are ongoing efforts to address this gap, including the introduction of the List of Essential In Vitro Diagnostics (EDL) and List of Priority Medical Devices for Cancer Management (PMDL), but much remains to be done, which is a key goal of our study.

## RESULTS

### The computational pipeline of DeepPT and ENLIGHT

#### Building cancer-type specific DeepPT models and their architecture

For each cancer type, a specific DeepPT model is constructed by training on formalin-fixed, paraffin-embedded (FFPE) whole slide images and their corresponding bulk gene expression profiles from TCGA patient samples. The model obtained can then be used to predict gene expression from new WSI both for internal held-out and external datasets.

In difference from previous studies aimed at predicting gene expression from WSI, which have focused on fine tuning the last layer of pre-trained convolutional neural networks (CNN), DeepPT is composed of three main components (**Methods,** **Figure 1a, and Supplementary Figure S1**): a CNN model for feature extraction, an auto-encoder for feature compression, and a multi-layer perceptron (MLP) for the final regression. Rather than training a model for each gene separately or for all genes together as was done in previous studies [39,41], we trained simultaneously on tranches of genes with similar median gene expression values, allowing shared signals to be leveraged while preventing the model from focusing on only the most highly expressed genes.

**Figure 1.**
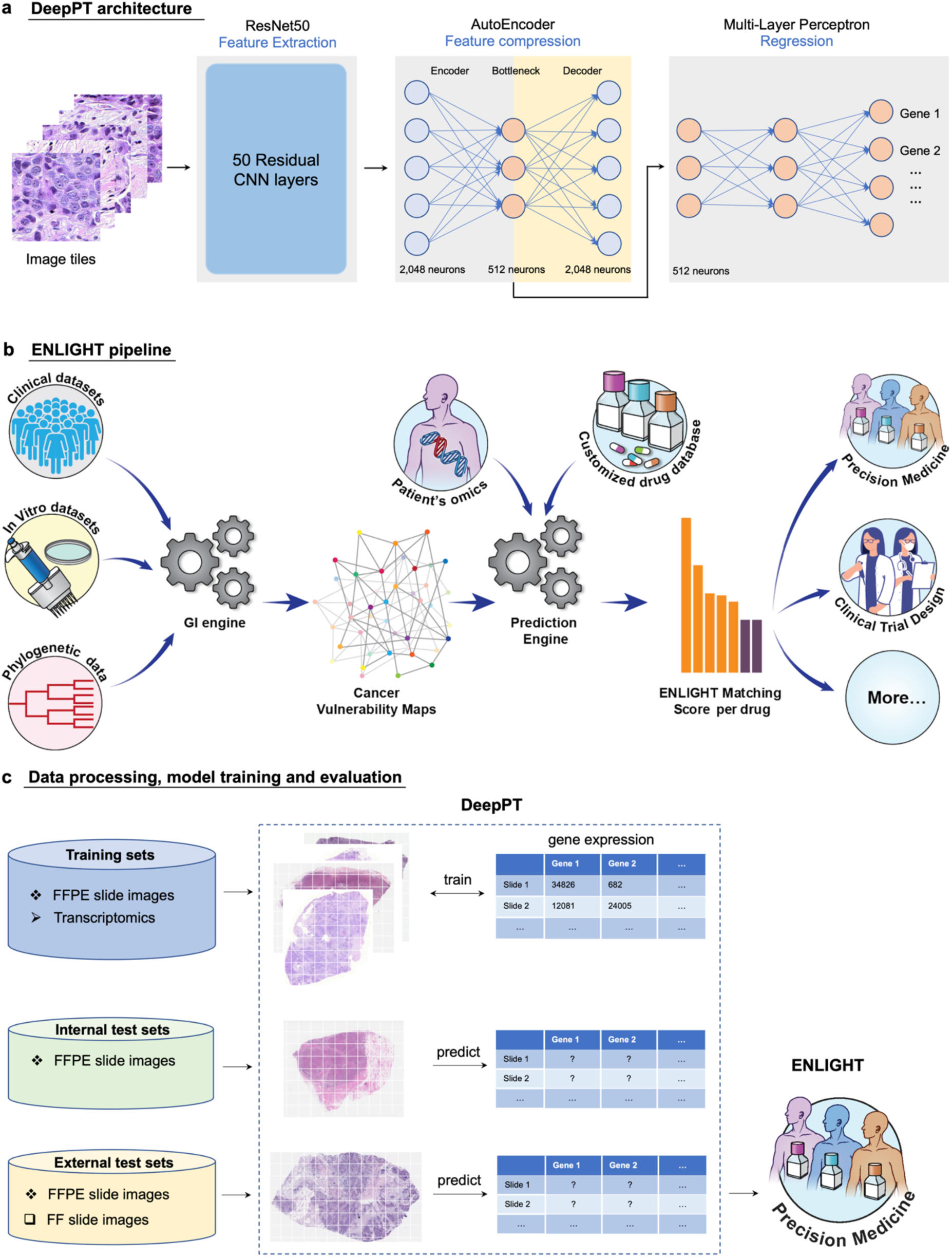
Study overview. (a) The three main components of DeepPT architecture, from left to right. The pre-trained ResNet50 CNN unit extracts histopathology features from tile images. The autoencoder compresses the 2,048 features to a lower dimension of 512 features. The multi-layer perceptron integrates these histopathology features to predict the sample’s gene expression. **(b) An overview of the ENLIGHT pipeline** (illustration taken from [44]: ENLIGHT starts by inferring the genetic interaction partners of a given drug from various cancer in-vitro and clinical data sources. Given the SL and SR partners and the transcriptomics for a given patient sample, ENLIGHT computes a drug matching score that is used to predict the patient response. Here, ENLIGHT uses DeepPT predicted expression to produce drug matching scores for each patient studied. **(c) Overview of the Analysis employing DeepPT and ENLIGHT: (i) top row:** DeepPT was trained with formalin-fixed paraffin-embedded (FFPE) slide images and matched transcriptomics for an array of different cancer types from the TCGA. **(ii) Middle row:** After the training phase, the models were applied to predict gene expression on the internal (held-out) TCGA datasets and on two external datasets on which they were never trained. **(iii) Bottom row:** The predicted tumor transcriptomics in each five independent test clinical datasets serves as input to ENLIGHT for predicting the patients’ response to treatment and assessing the overall prediction accuracy.

#### Predicting patient response from DeepPT-predicted expression using ENLIGHT – an overview

The predicted expression then serves as input to ENLIGHT [44], which is an algorithm that predicts individual responses to a wide range of targeted and immunotherapies based on gene expression measured from the tumor tissue (**Figure 1b**). ENLIGHT relies on the approach of analyzing functional genetic interactions (GI) around the target genes of a given therapy, originally presented in SELECT [46]. Specifically, two broad types of interactions are considered: Synthetic Lethality (SL), whereby the simultaneous loss of two non-essential genes is detrimental to the cell, and Synthetic Rescue (SR), whereby the loss of an essential gene can be compensated for through the over- or under-expression of a second “rescuer” gene. Based on the patient’s transcriptomics (whether measured from the tumor or predicted by DeepPT), ENLIGHT generates an ENLIGHT Matching Score (EMS), which evaluates the match between a patient and a treatment, (**Figure 1b**, [44], and **Methods** for more details).

#### Study design

The study design is depicted in **Figure 1c**. First, to build cancer type specific DeepPT models, we collected FFPE WSI together with matched RNAseq gene expression profiles for 16 cancer types from TCGA, composing 10 broad classes (some broad cancer indications include a few types, as per the original TCGA nomenclature): breast (BRCA), lung (LUAD and LUSC), brain (LGG and GBM), kidney (KIRC, KIRP and KICH), colorectal (COAD and READ), prostate (PRAD), gastric (STAD and ESCA), head and neck (HNSC), cervical (CESC), and pancreas (PAAD). These were chosen as they are major cancer types and/or have corresponding external datasets for evaluating purposes. Low quality slides (heavily marked, blurry, damaged, or too small) were excluded, resulting in 6,269 slides from 5,528 patients (**Table S1**). Each cancer type was processed, trained and evaluated separately. We performed a five-fold cross-validation to evaluate the model performance: In each loop, the patients were randomly split into five disjoint sets. Each of these sets was selected in turn as the *held-out test set* (20%), while the rest were used for training (80%). Note that the test set remained completely unseen during the model training, and the splits were performed at the patient level so that slides from the same patients are assigned to the same set to avoid information leakage between test and training sets.

Then, we performed external testing of DeepPT’s gene expression predictions. We applied the trained models of the respective cancer types to predict gene expression on two external datasets: the TransNEO breast cancer cohort (TransNEO-Breast) consisting of 160 fresh frozen (FF) slides [47], and an unpublished brain cancer cohort (NCI-Brain) consisting of 226 FFPE slides. Both datasets contained matched expression data (see **Methods**).

Our final goal is to use the inferred gene expression as input to ENLIGHT based predictions of patients treatment response. To this end, we further applied the DeepPT models to predict gene expression in five clinical trial datasets (see **Methods** and **Table S2** for full details). We show that combining ENLIGHT with DeepPT enables robust prediction of response to treatment in these independent test sets. Remarkably, this is done without adapting either DeepPT, which was trained once on TCGA samples, or ENLIGHT, which is not trained on any treatment response data in general, and specifically, on none of the clinical cohorts used here for evaluation purposes.

### Prediction of gene expression from histopathology images

As illustrated in **Figure 1**, we constructed models predicting normalized gene expression profiles from their corresponding histopathology images for each of 10 broad TCGA cancer classes. We then applied the trained models to predict gene expression of internal held-out test sets in each of the cancer types, using five-fold cross validation. In most cancer types, thousands of genes were significantly predicted, with Holm-Sidak corrected p-values < 0.05 (**Figure 2a** **and Supplementary Figure S2**). These results outperform the recently published state-of-the-art expression prediction approach, HE2RNA by Schmauch et al [41], in most cohorts considered in this study (**Figure 2a**). To further evaluate model performance, we estimated the Pearson correlation (R) between predicted and actual expression values of each gene across the test dataset samples. In most cancer types, thousands of genes had a correlation above 0.4 (**Supplementary Figure S3**). For breast cancer, for instance, DeepPT predicted 1,812 genes with mean correlation greater than 0.4, more than doubling the number of genes predicted at that correlation level reported by HE2RNA, which was 786 genes [41], further testifying to the increased accuracy of DeepPT.

**Figure 2.**
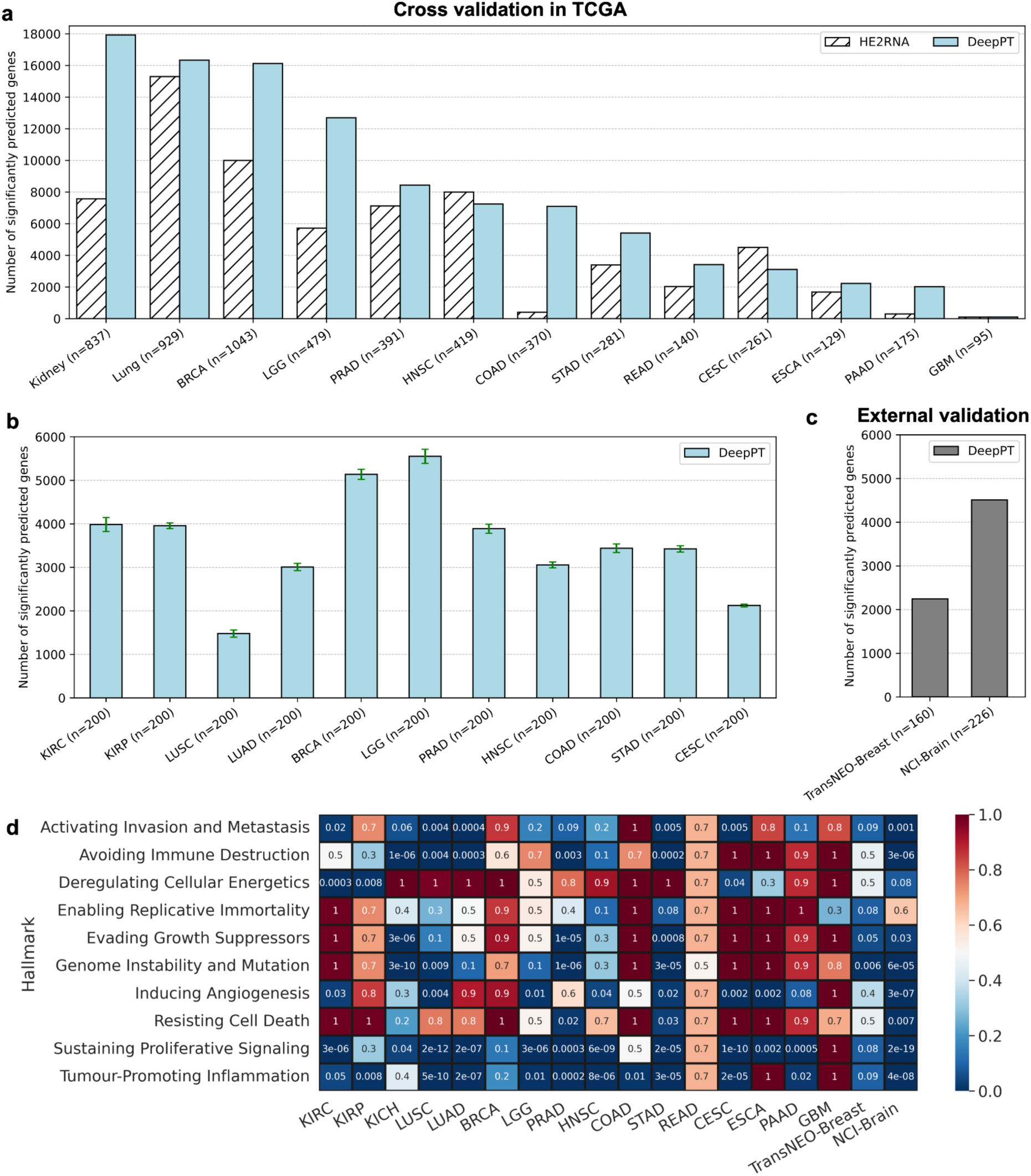
DeepPT prediction of gene expression from H&E slides. **(a) The number of significantly predicted genes for each TCGA cohort**, in comparison with the current state-of-the-art method, HE2RNA. For apples-to-apples comparison against HE2RNA, the performance of each cancer subtypes in Kidney (KIRC, KIRP, KICH) and Lung (LUSC, LUAD) are shown together, as reported in [41]. **(b) The number of significantly predicted genes, averaging over 30 randomly selected subsets.** Each subset comprises 200 samples that were randomly selected from the cohort. Only cohorts with at least 200 samples were analyzed. Error bars represent standard error of the mean. **(c) The number of significantly predicted genes in two independent test cohorts**, obtained by using pre-trained models on the corresponding TCGA cohorts. **(d) Pathway enrichment analysis on the significantly predicted genes.** Each row represents a different cancer hallmark and each column a different cohort (the two right columns correspond to the two external cohorts). Values denote the multiple hypothesis corrected p-value for pathway enrichment among the genes significantly predicted by DeepPT.

To assess the dependence of the number of significantly predicted genes on sample size, for each cohort, we randomly selected 30 subsets composed of 200 samples each (cohorts having less than 200 samples in total were not considered in this experiment) and measured the number of genes that were significantly predicted in each subsets. We observed that most cohorts had at least 3,000 genes that were significantly predicted (**Figure 2b**).

For external validation, we tested the prediction ability of DeepPT on two unseen independent datasets available to us, which contained matched tumor WSIs and gene expression. We first applied the DeepPT model constructed using the TCGA-Breast cancer dataset to predict gene expression from corresponding H&E slides of the TransNEO breast cancer cohort (n=160). Notably, the two datasets were generated independently at different facilities, with two different preparation methods (TCGA slides are FFPE while TransNEO slides are FF), so the histological features extracted from these two datasets are quite distinct (**Supplementary Figure S4**). Despite these differences, without any further training or tuning, we found 2,248 genes that were significantly predicted (**Figure 2c**). Similarly, we applied the DeepPT model trained on TCGA-Brain samples to predict gene expression from new unpublished NCI-Brain slides (n=226) and found that 4,510 genes were significantly predicted (**Figure 2c**). This testifies to the considerable predictive power and generalizability of DeepPT.

### Genes reliably predicted by DeepPT are enriched for cancer hallmarks

We next explored whether genes that are reliably predicted by DeepPT, i.e. those that are significantly correlated between predicted and measured expressions, have biological relevance to cancer. To this end, we carried out a pathway enrichment analysis (PEA) focused on cancer hallmarks. Specifically, we looked for enrichment among 10 cancer hallmarks described by Hanahan and Weinberg [48] and for which detailed gene sets were given by Iorio et al. [49]. **Figure 2d** summarizes the PEA results for all TCGA subtypes and the two external cohorts. Interestingly, we observed a strong enrichment for immune processes across the vast majority of cancer types (bottom row, “Tumor-promoting inflammation”), testifying to the important role of immune processes in shaping tumor morphology, as reflected by the slide images. Other enriched hallmarks include the cell cycle (“Sustaining proliferative signaling”), “Avoiding immune destruction” and “Activating invasion and metastasis”. Notably, these results are consistent and even stronger for the external datasets (TransNEO-Breast and NCI-Brain). They further testify that DeepPT can faithfully reconstruct key elements in cell expression related to cancer.

### DeepPT reconstructs prognostic signatures in TCGA

The observation that genes that promote proliferation and metastasis, which are well known prognostic markers, are specifically well predicted by DeepPT, led us to explore how well do such prognostic markers predict patient survival, when calculated over the gene expression predicted by DeepPT. To this end, we calculated three proliferation signatures known to be linked to cancer progression and poor prognosis using TCGA patients tumor data. These signatures include: (a) the expression of the MK67 gene, a well-known marker for cell proliferation, (b) the proliferation index derived in Whitfield et al. [50] and (c) an epithelial to mesenchymal transition (EMT) signature from MsigDB [51], associated with the formation and progression of metastasis. For each patient in each TCGA cohort, we calculated a signature score that is the mean gene-wise ranked expression across the genes of the signature. We then tested the correlation between signature scores derived from the predicted and the actual gene expression, and the association of each to patient survival using cox proportional hazard at the cohort level. All three signature scores exhibit a significant correlation between their actual and DeepPT-predicted expressions. Notably, the multi-gene EMT and proliferation signatures exhibit higher correlations (0.396 and 0.421) than the expression of the single MK67 gene (0.327), and a higher correlation than the individual genes that constitute these signatures (mean genewise correlation of 0.364 and 0.281 across each signature, respectively). Even though the correlation of the scores themselves are in the medium range, DeepPT reconstructs the prognostic value of the signatures fairly faithfully: the correlation between the hazard ratio of each signature between those computed based on the actual and predicted expressions is very high (0.77-0.88, **Figure 3**). These results testify that the ensemble of multiple genes yields a higher correlation when combined, as expected, and remarkably, the prognostic value of these signatures is well retained by DeepPT.

**Figure 3.**
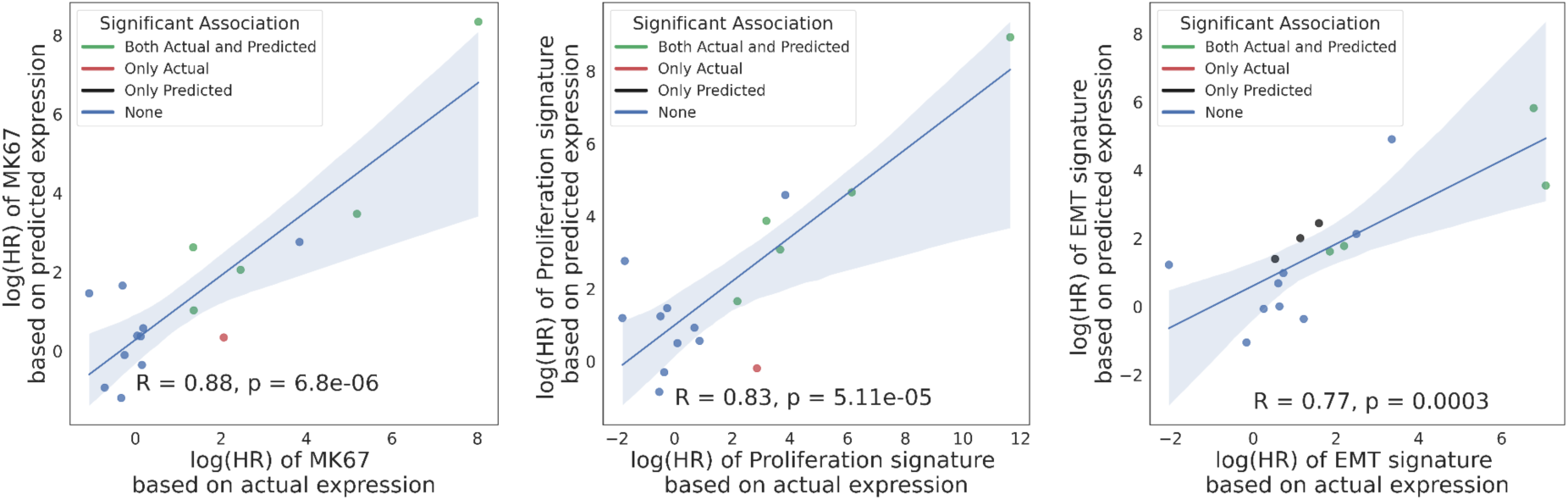
Comparison of the correlation of survival association in terms of log(HR) for three proliferation signatures (left: MK67; middle: Proliferation index; right: EMT pathway) based on actual (X axis) and predicted expressions (Y axis). Each point represents a different TCGA cohort, and points are color-coded according to the significance of survival association using a corrected p < 0.05 cutoff: green denotes that the survival association was significant by both the actual and predicted signatures, red/black only by the actual/predicted signatures, respectively. Pearson R and corresponding p-values are denoted in each panel.

### Predicting treatment response from DeepPT-imputed gene expression

As described earlier, the goal of ENLIGHT-DeepPT is to predict patients’ response from WSI without any training on the evaluation cohort. Given a previously unseen tumor slide image, we first apply the pre-trained, cancer-type specific DeepPT model to predict the tumor transcriptomics. Second, based on this predicted gene expression, we apply our published precision oncology algorithm, ENLIGHT [44], to predict the patient’s response.

We tested the ability of ENLIGHT-DeepPT to accurately predict patient response in five clinical cohorts, treated with various targeted and immunotherapies, for which patient slides and response data were available. Those include two HER2+ breast cancer patient cohorts treated with chemotherapy plus Trastuzumab [47,52], a BRCA+ pancreatic cancer cohort treated with PARP inhibitors (Olaparib or Veliparib), a mixed indication cohort of Lung, Cervical and Head & Neck patients treated with Bintrafusp alfa [53], a bi-specific antibody that targets TGFB and PDL1, and finally, an ALK+ NSCLC cohort treated with ALK inhibitors (Alectinib or Crizotinib). For each dataset, the response definition was determined by the clinicians running the respective trial (see **Methods** and **Table S2** for more details). As ENLIGHT does not predict response to chemotherapies, only the targeted agents were considered for response prediction.

For each cohort, we used the DeepPT model previously trained on the appropriate TCGA cohort, *without any changes and with no further training*, to predict the gene expression values from the H&E slide of each patient’s pre-treated tumor. We then applied ENLIGHT to these predicted gene expression values to produce ENLIGHT Matching Scores (EMS) based on the genetic interaction network of the given drug, as was originally published in [44]. Importantly, we do not restrict the GI network to include only genes with strong correlations between the actual and predicted expression values; This is done as ENLIGHT considers the combined effect of a large set of genes, averaging out noise arising from individual gene expression prediction. Notably, restricting ENLIGHT’s GI networks to include only significantly predicted genes does not improve results (**Supplementary Figure S9)**.

The prediction accuracy of ENLIGHT-DeepPT in each of these five datasets individually and in aggregate is shown in **Figure 4**. Since the ENLIGHT-DeepPT workflow is designed with clinical applications in mind, we focus our assessment of its predictive power on measures that have direct clinical importance, including both the odds ratio (OR) of response and the average precision. The OR denotes the ratio of the odds to respond among patients receiving ENLIGHT-matched treatments vs. the odds to respond among patients whose treatments were not ENLIGHT-matched. Patients were considered ENLIGHT-matched if their EMS scores were greater or equal to a threshold value of 0.54. This threshold was determined already in the original ENLIGHT publication [44] on independent data and was kept fixed here. Using this predefined threshold, we observe that the OR of ENLIGHT-DeepPT is higher than 1 for all datasets, though this was not statistically significant for the PARPi and ALKi datasets, probably due to their small sample sizes (**Figure 4a**). This demonstrates that patients receiving ENLIGHT-matched treatments indeed had a higher chance to respond.

**Figure 4.**
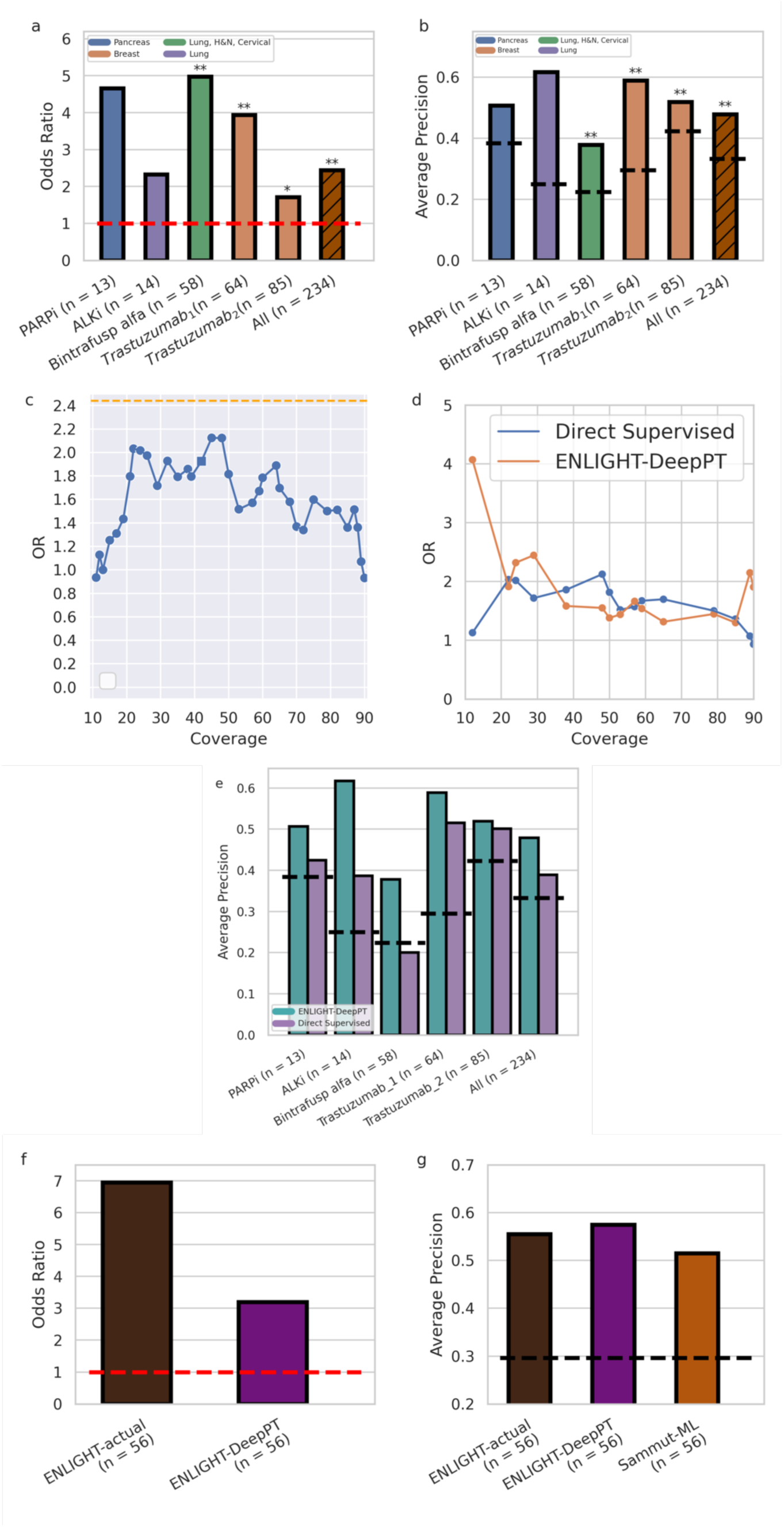
Predicting treatment response from H&E slides. **(a)** Odds Ratio (OR, Y axis) for the five datasets tested and the aggregate cohort of all patients together (X axis). Drug and sample sizes are denoted in the X axis labels. Orange horizontal dashed line denotes an OR of 1 which is expected by chance. Bars are color coded according to the indication(s) of the respective cohort. Asterisks denote significance of OR being larger than 1 according to Fisher’s exact test **(b)** Average Precision (AP, Y axis) for the five datasets and the aggregate cohort, as in **a**. Black horizontal dashed lines denote the ORR for each dataset. An AP higher than the ORR demonstrates better accuracy than expected by chance. Asterisks denote significance of AP being higher than response rate using one-sided proportion test. (**c**) OR of the Direct Supervised method (Y axis) for all 234 patients as a function of the fraction of patients above a given threshold (coverage, X axis). We present only coverage between 10-90% to avoid the measurement noise of extreme coverage values, where data is too small. Orange dashed line denotes the OR of ENLIGHT-DeepPT for all 234 patients at its original clinical decision threshold. The square denotes the threshold on the Direct Supervised that yields the same coverage as ENLIGHT-DeepPT at its original, fixed threshold. (**d**) Comparison of the OR of ENLIGHT-DeepPT and the Direct Supervised methods (Y axis) at thresholds that yield the same coverage (X axis). (**e**) Average Precision of ENLIGHT-DeepPT (cyan) and Direct Supervised (purple) for each dataset and on aggregate as in **b**. Dashed lines denote the ORR for each case as in **b (f)** OR for ENLIGHT-actual and ENLIGHT-DeepPT when predicting response to Trastuzumab (for the Trastuzumab_1_ cohort). **(g)** Comparison of AP (Y axis) for both ENLIGHT based models and the Sammut-ML predictor of Sammut et al. [47]. All methods were applied to the same patient group. Black horizontal dashed line denotes the ORR. All p-values were FDR corrected. * = p < 0.1, ** = p < 0.05.

Complementing the OR measure, which is of major translational interest as it quantifies the performance at a specific decision threshold, **Figure 4b** further depicts the prediction performance of ENLIGHT-DeepPT via a complementary measure, the average precision (AP). AP is a measure of the precision of a prediction model across the entire range of thresholds. For a classifier to be of merit, its AP should be higher than the overall response rate (ORR), which denotes the fraction of responders observed in each cohort. Reassuringly, the AP of ENLIGHT-DeepPT well exceeds the ORR for all five datasets, testifying to its broader predictive power beyond that quantified by the OR.

Turning to an aggregate analysis of the performance of ENLIGHT-DeepPT, by analysing all patients together in a simulated “basket” trial in which each patient receives a different treatment (n = 234), the OR of ENLIGHT-DeepPT is 2.44 ([1.36,4.38] 95% CI, left bar in **Figure 4a**), significantly higher than 1 (p = 0.002, Fisher’s exact test), and its precision is 47.8%, a 43.5% increase compared to the overall response rate of 33.3% (*p* = 1.28×10^−6^, one sided proportion test). The AP is 0.48 (left bar, **Figure 4b**), significantly higher than the baseline response rate (p = 0.0003, one sided permutation test).

One of the advantages of ENLIGHT’s unsupervised approach is that it does not rely on data labelled with response to treatment data which is usually scarce. Such data is required for training supervised models, which in theory, given sufficiently large datasets, are expected to yield higher performance on unseen datasets than unsupervised methods like ENLIGHT. The question remains whether such supervised methods are advantageous for realistically small datasets as studied in this work. To compare the performance of our two-step indirect model with a direct supervised model, we trained the same computational deep learning pipeline as the one used in DeepPT on H&E slides and their corresponding response data on each of the five evaluation datasets described above, except that we replaced the regression component with a classification component. We used the same training strategy that has been widely applied previously in the literature for direct H&E slide-based classification [6,9,20,23,42] (see **Methods** for details), and termed this the *Direct Supervised* model. Due to the lack of independent treatment data for training, we applied leave-one-out cross-validation (LOOCV) to evaluate the performance of the direct supervised models. This in-cohort training gives an inherent advantage to the Direct Supervised approach over ENLIGHT-DeepPT, which was not exposed to these datasets at all. Since no tuning data was available to calibrate a single threshold for the Direct Supervised methods as was done in [44], we calculated the OR of the Direct Supervised for all patients (n = 234) on all possible thresholds. **Figure 4c** presents the OR as a function of the coverage (the fraction of patients with scores above a specific threshold). We compared these values to the OR of ENLIGHT-DeepPT at the clinical decision threshold established in [44] based on measured RNA levels (0.54, red dashed line). Surprisingly, no threshold on the Direct Supervised values yields an OR that surpasses the OR of ENLIGHT-DeepPT at its predetermined clinical decision threshold. In addition, we calculated ENLIGHT-DeepPT’s OR at various possible EMS thresholds and compared the results at thresholds that yield the same coverage in both ENLIGHT-DeepPT and Direct Supervised (**Figure 4d**). Finally, **Figure 4e** compares the average precision (which is threshold independent) of the two models. Remarkably, the overall performance of ENLIGHT-DeepPT, being an unsupervised method, as measured by OR and AP, is comparable with that of the supervised classifiers trained and predictive only for specific treatments, and in some cases ENLIGHT-DeepPT even outperforms them. Moreover, training in-cohort, as was done here for the supervised methods due to lack of sufficient data, has a clear risk of overfitting.

Ideally, an external dataset is required to study the generalizability of the model. Among the datasets of this study, this was possible only for Trastuzumab which appeared in more than one dataset. When we tested the model trained on one Trastuzumab dataset on the other set and vice versa we saw low generalizability: the AP went down from 0.52 in LOOCV for Trastuzumab_1_ to 0.27 when this model was tested on Trastuzumab_2_, and from 0.5 to 0.43 in the other direction. This suggests that the results obtained for the supervised models may be overfitted. Clearly, for cases where large data exists, a supervised method can outperform ENLIGHT-DeepPT. In contrast, ENLIGHT-DeepPT is unsupervised, and the results presented here testify to its potential generalizability. In addition, any supervised model can only be obtained for drugs with coupled H&E and response data (4 drugs in this study), while ENLIGHT-DeepPT can produce predictions to virtually any targeted treatment.

For one of the datasets (Trastuzumab_1_), RNA sequencing of the tumor gene expression was also available and was previously analyzed by ENLIGHT in [44]. **Figures 4f and 4g** compare the predictive performance using ENLIGHT-DeepPT scores to that using ENLIGHT scores calculated based on the *measured* expression values (denoted *ENLIGHT-actual*). Using the previously established threshold of 0.54, the OR of ENLIGHT-DeepPT is 3.2, which is lower than the OR of 6.95 obtained by ENLIGHT-actual, but still significantly higher than expected by chance (p = 0.02, test for OR > 1). The positive predictive value (PPV) (also known as precision) of ENLIGHT-DeepPT was 53.3%, slightly higher than but not significantly different from the PPV of 52% when using ENLIGHT-actual, and 80% higher than the basic ORR of 29.7% observed in this study. However, the sensitivity (the fraction of responders correctly identified) of ENLIGHT-DeepPT is markedly lower than that of ENLIGHT-actual, 42.1% vs. 68.%.

Finally, we sought to compare ENLIGHT-DeepPT to other predictive models for drug response. For the drugs analyzed in this study, the only available mRNA-based model for response is the multi-omic machine learning predictor that uses DNA, RNA and clinical data, published by Sammut et al. [47] denoted here as Sammut-ML. This model was based on *in-cohort supervised learning* to predict response to chemotherapy with or without trastuzumab among HER2+ breast cancer patients. **Figure 4g** compares ENLIGHT-actual and ENLIGHT-DeepPT performance to Sammut ML. In both analyses, we applied all methods to the same patient group of 56 patients for whom all relevant data was available (RNAseq, H&E slide, DNAseq and clinical features). To systematically compare between the predictors across a wide range of decision thresholds, and since Sammut et al. did not derive a binary classification threshold, we used AP as the comparative rod here. As can be seen, all methods have quite comparable predictive power, with ENLIGHT-DeepPT having the highest AP (difference not statistically significant). Importantly, using only H&E slides without need for RNA or DNA data or other clinical features has an invaluable practical advantage. Notably, the predictions for Trastuzumab_1_ were made on fresh frozen tissue slides, which differ considerably from FFPE samples used to train the DeepPT model, testifying to the robustness of DeepPT and ENLIGHT. To complement this analysis, we show that ENLIGHT-DeepPT outperforms a model that uses only the predicted expression of the drug targets as predictors of response (**Supplementary Figure S10**).

## DISCUSSION

Our study demonstrates that combining DeepPT, a novel deep learning framework for predicting gene expression from H&E slides, with ENLIGHT, a published unsupervised computational approach for predicting patient response from pre-treated tumor transcriptomics, could be used to form a new *ENLIGHT-DeepPT* approach for H&E-based prediction of clinical response to a host of targeted and immune therapies. We began by showing that DeepPT significantly outperforms the current state-of-the-art method in predicting mRNA expression profiles from H&E slides. Then, we showed that the aggregate signal from multiple genes can overcome weak correlation at the individual gene level. Finally, and most importantly, *ENLIGHT-DeepPT* successfully predicts the true responders in several clinical datasets from different indications, treated with a variety of targeted drugs directly from the H&E images, demonstrating its potential clinical utility throughout. Notably, its prediction accuracy on these datasets is on par with that of direct predictors from the images, as is the current practice, even though the latter have been trained and tested in a cross validation in-cohort manner.

Combining DeepPT with ENLIGHT is a promising approach for predicting response directly from H&E slides because it does not require response data on which to train. This is a crucial advantage compared to the more common practice of using response data to train classifiers in a supervised manner. Indeed, while sources like TCGA lack response data that would enable building a supervised predictor of response to targeted and immune treatments, applying ENLIGHT to predicted expression has successfully enabled the prediction of response to four different treatments in five datasets spanning six cancer types with considerable accuracy, without the need for any treatment data for training. While supervised models can only be obtained for drugs with available H&E and response data, ENLIGHT-DeepPT can produce predictions to virtually any targeted treatment, and importantly, including ones in early stages of development where such training data is still absent.

DeepPT is fundamentally different from previous computational pipelines for gene expression prediction both in its model architecture and in its training strategy. We attribute its superior performance to four key innovations: (i) All previous studies fed output from the conventional pre-trained CNN model (trained with natural images from ImageNet database) directly into their regression module, whereas we added an auto-encoder to re-train the output of the pre-trained CNN model. This helps to exclude noise, avoid overfitting and reduce the computational demands. (ii) Our regression module is an MLP model in which the weights from the input layer to the hidden layer are shared among genes. This architecture enables the model to exploit the correlations between the expression of the genes. (iii) We trained together sets of genes with similar median gene expression values; doing so further implements a form of multitask learning and prevents the model from focusing on only the most highly expressed genes. (iv) We performed ensemble learning by taking the mean predictions across all models. This further improves the prediction accuracy quite significantly (**Supplementary Figure S6**).

DeepPT can be broadly applied to all cancer types for which H&E slides and coupled mRNA data are available for training; however, similar to many other deep learning models, it requires a considerable number of training samples comprising matched imaging and gene expression. An interesting direction for future work would be to apply transfer learning between cohorts, to improve the predictive performance in cancer types with yet small training cohorts. In other words, it might be possible to train the model on large TCGA cohorts such as breast and lung cancer, then fine tune it for generating predictions for cancer types with smaller TCGA cohorts such as melanoma or ovarian cancer.

A notable finding of this study is the robustness of response predictions based on H&E slides when combining DeepPT and ENLIGHT. First, despite the inevitable noise introduced by the prediction of gene expression, the original ENLIGHT GI networks, designed to predict response from measured RNA expression, worked well as-is in predicting response based on the DeepPT-predicted expression. In fact, when restricting the GI networks to include only significantly predicted genes, the results are not improved. Second, though DeepPT was trained using FFPE slides, it generalized well and could be used as-is to predict expression values from FF slides. This demonstrates the applicability of DeepPT for predicting RNA expression either from FF or from FFPE slides. Nevertheless, as promising as the results presented here are, they should of course be further tested and expanded upon by applying the generic pipeline presented here to many more cancer types and treatments.

There are several limitations of our study that should be noted: (i) as explained above, while DeepPT can predict expression reliably for many genes based on a relatively small number of samples, it is still reliant on sufficient training data and future studies should aim to enlarge the number of genes whose expression is accurately predicted. (ii) ENLIGHT-DeepPT should be more extensively validated on additional indications and treatments, as data is accumulated. (iii) Here we used ENLIGHT’s clinical decision threshold as is from [44]. Further research is required to fine tune the decision threshold for ENLIGHT-DeepPT once its application to a broader set of cohorts becomes feasible.

Developing a response prediction pipeline from H&E slides, if reasonably accurate and further carefully tested and validated in clinical settings, could obviously be of utmost benefit, as NGS results often take 4-6 weeks after initiation to return a result. Many patients who have advanced cancers require treatment immediately, and this method can potentially offer treatment options within a shorter time frame. Moreover, obtaining H&E images can be done at relatively low cost, compared to the expenses incurred by NGS. Increasing efforts to harness the rapid advances in deep learning are likely to improve precision oncology approaches, including by leveraging histopathology images. Given its general and unsupervised nature, we are hopeful that *ENLIGHT-DeepPT* may have considerable impact, making precision oncology more accessible to patients in Low- and middle-income countries (LMICs), in under-served regions and in other situations where sequencing is less feasible. Specifically, affordable cancer diagnostics is critical in LMICs since their limited access to cancer diagnostics is a bottleneck for effectively leveraging the increasing access to cancer medicine [45]. While promising, one should of course cautiously note that the results presented in this study await a broader testing and validation in carefully designed *prospective studies* before they may be applied in the clinic. We are hopeful that the results presented here will expedite such efforts by others going forward.

## METHODS

### Data collection

- TCGA histological images and their corresponding gene expression profiles were downloaded from GDC (https://portal.gdc.cancer.gov). Only diagnostic slides from primary tumors were selected, making a total of 6,269 formalin-fixed paraffin-embedded (FFPE) slides from 5,528 patients with breast cancer (1,106 slides; 1,043 patients), lung cancer (1,018 slides; 927 patients), brain cancer (1,015 slides; 574 patients), kidney cancer (859 slides; 836 patients), colorectal (514 slides; 510 patients), prostate (438 slides; 392 patients), gastric (433 slides; 410 patients), head & neck (430 slides; 409 patients), cervical (261 slides; 252 patients), pancreatic cancer (195 slides; 175 patients).

- The TransNEO-Breast dataset consists of fresh frozen slides and their corresponding gene expression profiles from 160 breast cancer patients. Full details of the RNA library preparation and sequencing protocols, as well as digitisation of slides have been previously described [47].

- The NCI-Brain histological images and their corresponding gene expression profiles were obtained from archives of the Laboratory of Pathology at the NCI, and consisted of 226 cases comprising a variety of CNS tumors, including both common and rare tumor types. All cases were subject to methylation profiling to evaluate the diagnosis, as well as RNA-sequencing.

- The Bintrafusp alfa treated cohort consisted of 58 patients with lung cancer (9 patients), cervical cancer (16 patients), and head and neck cancer (33 patients). FFPE slides were made available from the NCI.

- The Trastuzumab_1_ cohort is a subset of TransNEO-Breast dataset mentioned above, consisting of 64 patients who had received a combination of chemotherapy and Trastuzumab.

- Trastuzumab_2_, a HER2+ breast cancer cohort treated with a combination of Trastuzumab and chemotherapy, consisted of 85 patients and their FFPE slides [36, 50]. FFPE slides were downloaded from The Cancer Imaging Archive database (TCIA, https://www.cancerimagingarchive.net).

- The ALKi dataset consisted of 14 NSCLC ALK mutated patients treated with Crizotinib or Alectinib. Corresponding FFPE slides were made available from Colorado University.

- The PARPi dataset consisted of 13 germline BRCA mutated pancreatic cancer patients treated with PARP inhibitors. FFPE slides were made available from Sheba Medical Center.

For each dataset, the classification of patients to responders and non-responders was based on the criterion used by the clinicians running the respective trial: for Trastuzumab_1_ and Trastuzumab_2_, response was defined as pCR while non-responders were defined as RD. For Bintrafusp alfa, responders were defined as patients with partial response or complete response. For ALKi, response was defined as more than 18 months progression free survival. For PARPi, response was defined as more than 36 months overall survival. Full details can be found in **Table S2**.

### Histopathology image processing

We first used Sobel edge detection [54] to identify areas containing tissue within each slide. Because the WSI are too large (from 10,000 to 100,00 pixels in each dimension) to feed directly into the deep neural networks, we partitioned the WSI at 20x magnification into non-overlapping tiles of 512 x 512 RGB pixels. Tiles containing more than half of the pixels with a weighted gradient magnitude smaller than a certain threshold (varying from 10 to 20, depending on image quality) were removed. Depending on the size of slides, the number of tiles per slide in the TCGA cohort varied from 100 to 8,000 (**Supplementary Figure S7**). In contrast, TransNEO slides for example are much smaller, resulting in 100 to 1,000 tiles per slide (**Supplementary Figure S7e**). To minimize staining variation (heterogeneity and batch effects), color normalization was applied to the selected tiles.

### Gene expression processing

Gene expression profiles were obtained as read counts for approximately 60,483 gene identifiers. Genes considered expressed were identified using edgeR, resulting in approximately 18,000 genes for each cancer type. The median expression across samples of each gene varied from 10 to 10,000 reads for each dataset (**Supplementary Figure S8**). To reduce the range of gene expression values, and to minimize discrepancies in library size between experiments and batches, a normalization was performed as described in our previous work [44].

### DeepPT architecture

Our model architecture was composed of three main units (**Supplementary Figure 1**).

<u>(1) Feature extraction:</u> The pre-trained ResNet50 CNN model, trained with 14 million natural images from the ImageNet database [55,56] was used to extract features from image tiles. Before feeding these tiles into the ResNet50 unit, the image tiles were resized to 224 x 224 pixels to match the standard input size for the convolutional neural network. Through the feature extraction process, each input tile is represented by a vector of 2,048 derived features.

<u>(2) Feature compression:</u> We applied an autoencoder, which consists of a bottleneck of 512 neurons, to reduce the number of features from 2,048 to 512. This helps to exclude noise, to avoid overfitting, and finally to reduce the computational demands. As shown in **Supplementary Figure S5**, a large number of ResNet features are constantly zero (**Supplementary Figure S5**, upper panels). This data sparsity is considerably reduced in the autoencoder features (**Supplementary Figure S5**, lower panels).

<u>(3) Multi-Layer Perceptron (MLP) regression:</u> The purpose of this component is to build a predictive model linking the aforementioned auto-encoded features to whole-genome gene expression. The model consists of three layers: (1) an input layer with 512 nodes, reflecting the size of the auto-encoded vector; (2) a hidden layer whose size depends on the number of genes under shared consideration; and (3) an output layer with one node per gene. The rationale behind this architecture is to leverage similarity among the genes under shared consideration, as captured by the weights connecting the input layer to the hidden layer. The weights connecting the hidden layer to the output layer model the subsequent relationship between the hidden layer and each individual gene. This follows the philosophy of multi-task learning. If the prediction of each gene’s expression level represents a single task, then our strategy is to first group these tasks for shared learning, followed by optimization of each individual task. In our default whole-genome approach, we bin genes into groups of 4,096 whose median expression levels are similar, and we use 512 hidden nodes. Because the training data consists of gene expression at the slide level (i.e. bulk gene expression, as opposed to at spatial resolution), we average our per-tile predictions to obtain a mean value at the slide level.

### DeepPT training and evaluation

We trained and evaluated each cancer type independently. To evaluate our model performance, we applied 5x5 nested cross-validation. For each outer loop, we split the entire patients (of each cohort) into training (80%) and held-out test (20%) set. We further split the training set into internal training and evaluation set, according to five-fold cross validation. The models were trained and evaluated independently with each pair of training/validation sets. Averaging the predictions from the five different models represents our final prediction for each single gene on each held-out test set. We repeated this procedure five times across the five held-out test sets, making a total of 25 trained models. These models trained with TCGA cohorts were used to predict the expression of each gene in a given external cohort by computing the mean over the predicted values of all models. Because each patient can have more than one slide, we average the slide-level predictions to obtain patient-level predictions.

As noted in the Model Architecture section, tranches of genes with similar median expression levels were grouped for simultaneous training and evaluation. This was done to optimize model performance and model efficiency, and contrasts with approaches in the literature that either train on each gene separately [39] or on all genes together [41]. Each training round was stopped at a maximum of 500 epochs, or sooner if the average correlation per gene between actual and prediction values of gene expression on the validation set did not improve for 50 continuous epochs. The Adam optimizer with mean squared error loss function was employed in both auto-encoder and MLP models. A learning rate of 10^−4^ and a minibatches of 32 image tiles per step were used for both the auto-encoder model and MLP regression model. To reduce overfitting, a dropout of 0.2 was used.

### Data augmentation

Because the number of samples in the TCGA-PAAD cohort is relatively small, to further reduce overfitting, we artificially increased the amount of data for this cohort by rotating the whole slide images by 90°, 180°, 270°. During the test time, the average of four symmetries represents our prediction for each slide. No data augmentation was performed for other cohorts due to its high computational demand.

### Direct supervised model

The direct (end-to-end) supervised model was designed to classify responders and non-responders directly from their slides, without the intermediate step of gene expression prediction. To this end, we applied the same computational deep learning framework that was used for prediction of gene expression, except that the MLP regression component was replaced by an MLP classification component. Each of the five evaluation datasets were processed independently. Following previous approaches [6,9,20,23,42], all tiles from a given slide inherit the slide label. Because the number of samples for each dataset is relatively small, we evaluated the direct supervised models using leave one out cross-validation. For each held-out patient, we applied a bootstrap sampling technique to randomly split the remaining patients into training (80%) and validation (20%) sets 30 times, resulting in 30 models. For each model, slide-level prediction was computed by averaging tile-level predictions within that slide. The final prediction of each held-out slide was computed by averaging its predictions over the 30 models.

### Implementation details

All analysis in this study was performed in Python 3.9.7 and R 4.1.0 with the libraries including Numpy 1.20.3, Pandas 1.3.4, Scikit-learn 1.1.1, Matplotlib 3.4.3, and edgeR 3.28.0. Image processing including tile partitioning and color normalization was conducted with OpenSlide 1.1.2, OpenCV 4.5.4, PIL 8.4.0. The histopathology feature extraction was carried out using TensorFlow 2.8.0. The feature compression (autoencoder unit) and MLP regression parts were implemented using PyTorch 1.12.0. Pearson correlation was calculated using Scipy 1.5.0.

### ENLIGHT

ENLIGHT’s drug response prediction comprises two steps: (i) Given a drug, the *GI engine* identifies the clinically relevant genetic interaction (GI) partners of the drug’s target gene(s). The GI engine first identifies a list of initial candidate Synthetic lethal/Synthetic rescue (SL/SR) interactions by analyzing cancer cell line dependencies based on the principle that SL/SR interactions should decrease/increase tumor cell viability, respectively, when ‘activated’ (e.g., in the SL case, viability is decreased when both genes are under-expressed). It then selects those pairs that are more likely to be clinically relevant by analysing a database of tumor samples with associated transcriptomics and survival data, requiring a significant association between the joint inactivation of target and partner genes and better patient survival for SL interactions, and analogously for SR interactions. (ii) The drug-specific GI partners are then used to predict a patient’s response to each drug based on the gene expression profile of the patient’s tumor. The ENLIGHT Matching Score (EMS), which evaluates the match between patient and treatment, is based on the overall activation state of the set of GI partner genes of the drug targets, deduced from the gene expression, reflecting the notion that a tumor would be more susceptible to a drug that induces more active SL interactions and fewer active SR interactions.

We applied ENLIGHT in its original version as described in [44]. The only modification made to the original ENLIGHT version is the exclusion of the component that, in the case of monoclonal antibodies, considers the expression of the drug target itself. As including this component does not increase the prediction accuracy (**Supplementary Figure S11**), we excluded it as it highly weighs a single gene and is hence much more susceptible to perturbation resulting from noisy prediction of that one gene”.

## Data and Code Availability

TCGA data are available from https://portal.gdc.cancer.gov. TCIA data is available from https://www.cancerimagingarchive.net/. All DeepPT predicted expressions and relevant response data, along with the code to calculate performance measures are available in Github: https://github.com/PangeaResearch/enlight-deeppt-data. ENLIGHT scores given expression profiles (either measured directly from the tumor or predicted from slides) can be calculated using a web service at ems.pangeabiomed.com. The DeepPT prediction pipeline will be made accessible upon acceptance via https://zenodo.org/record/7912194.

## Acknowledgments

This work was supported by the Australian Research Council (ARC) (D.T.H, E.A.S) and by the Intramural Research Program of the National Institutes of Health (NIH), National Cancer Institute (NCI), Center for Cancer Research (CCR) (S.S, N.S, E.R). This work utilized the super computational resources of the Australian National Computational Infrastructure (AUNCI) and the Australian National University Merit Allocation Scheme (ANUMAS).

## SUPPLEMENTARY MATERIAL

**Figure S1.**
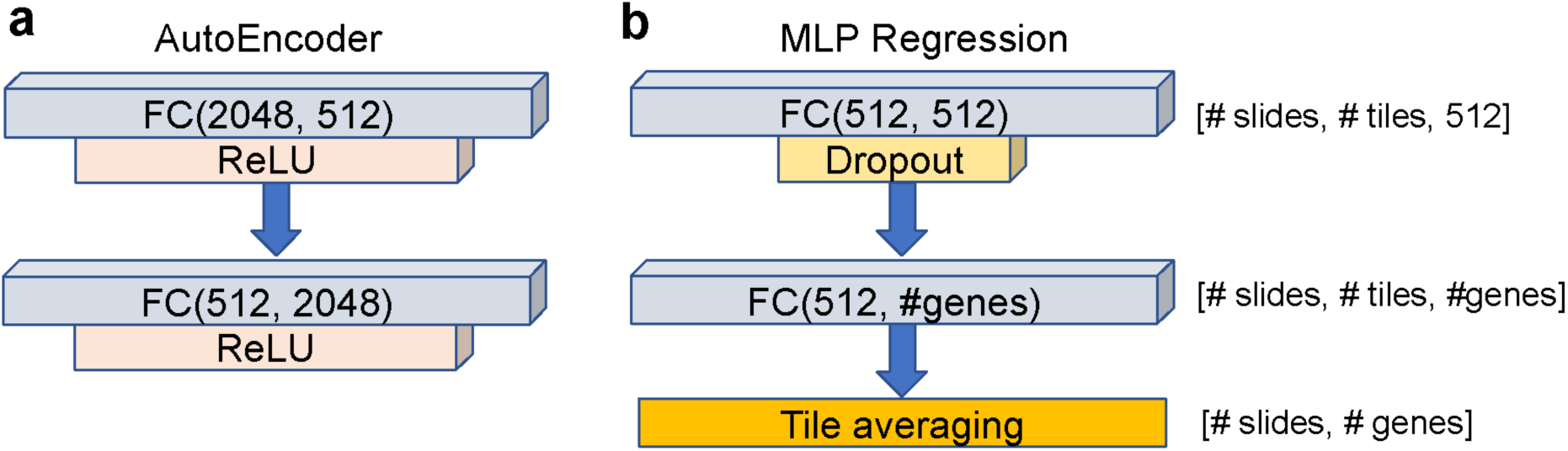
Model architecture in detail. (a) The feature compression subnetwork consists of an input layer of 2,048 neurons, a bottleneck of 512 neurons, and an output layer of 2,048 neurons. (b) The MLP regression subnetwork consists of an input layer of 512 neurons, a hidden layer of 512 neurons, and an output layer with the number of neurons reflecting the number of genes in each group.

**Figure S2.**
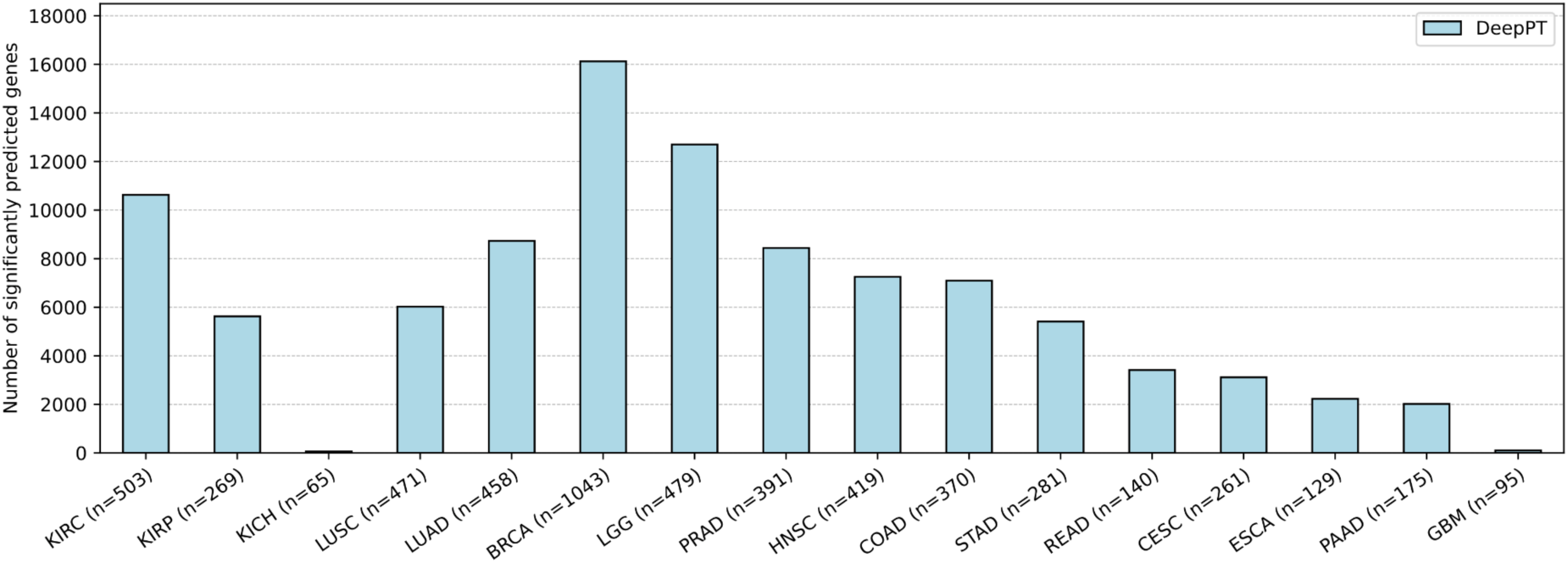
DeepPT performance. The number of genes that were significantly predicted by DeepPT in which the result for each TCGA cancer cohort was shown separately.

**Figure S3.**
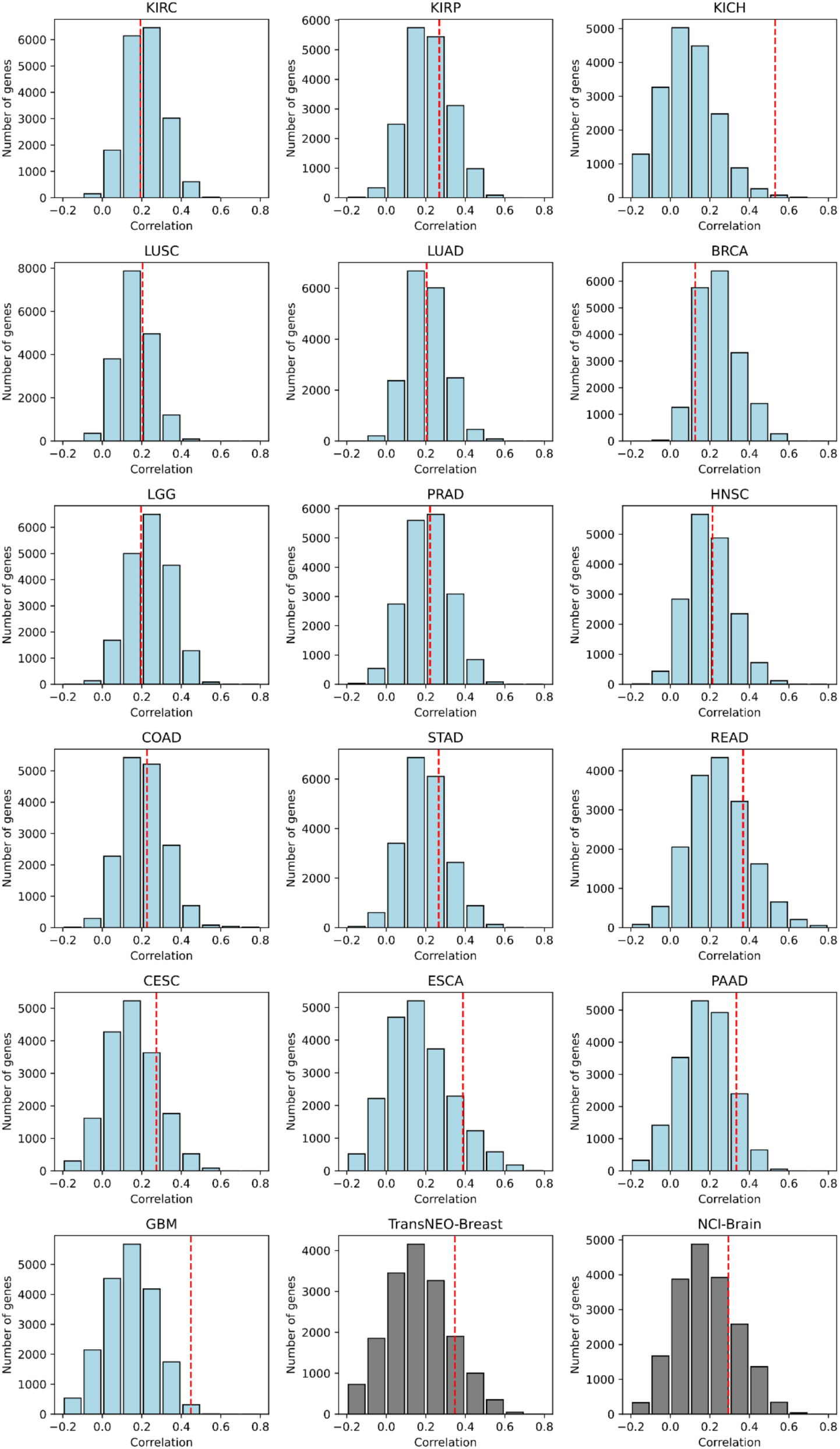
Histograms of the Pearson correlation coefficients between predicted and actual expression for each gene across test sets, for 16 TCGA cohorts (light blue) and 2 external cohorts (gray). Red dashed lines represent the correlation coefficient level beyond which the results are significant (p-value < 0.05 after correction for multiple hypotheses testing).

**Figure S4.**
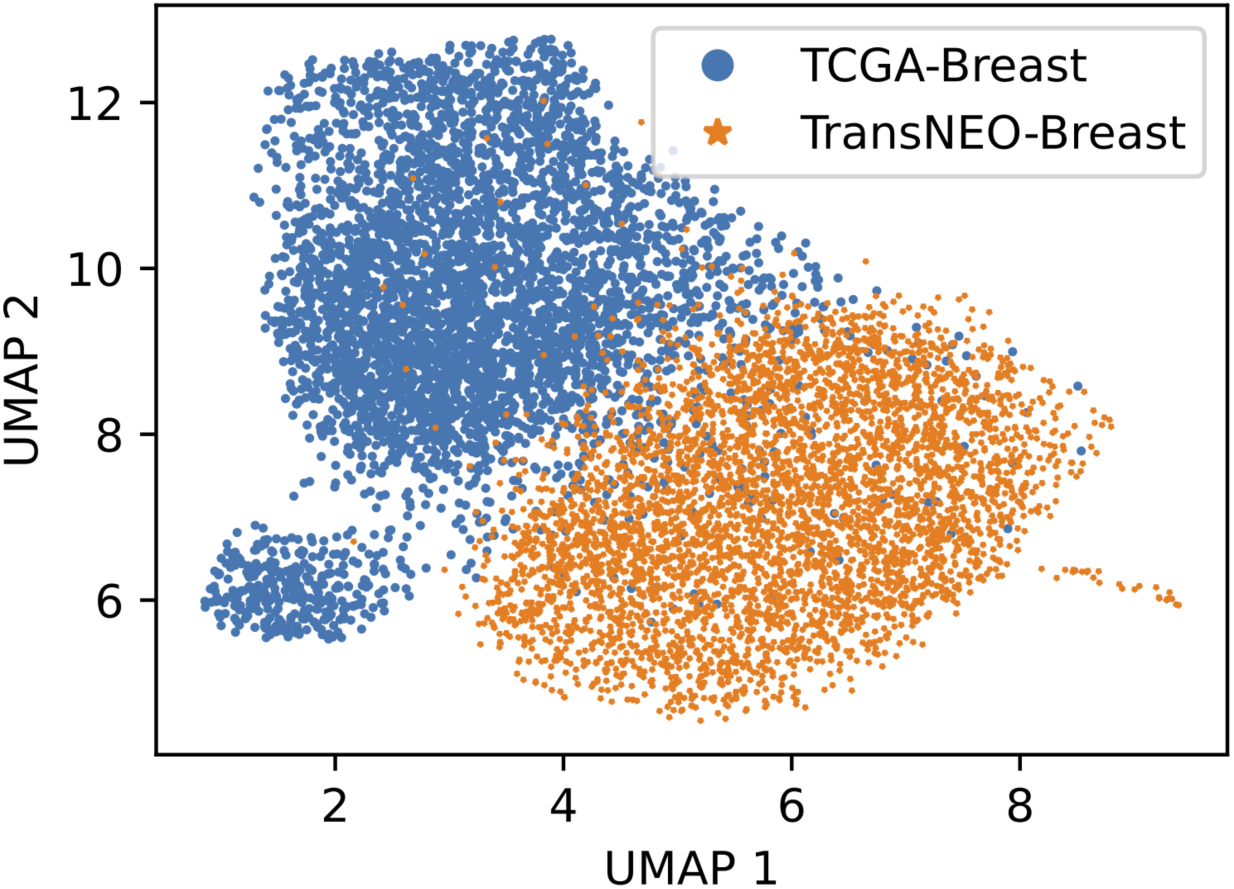
Difference between histopathological features extracted from TCGA-Breast tiles and TransNEO-Breast tiles. UMAP visualization of 2,048 histopathological features that were extracted by using pre-trained ResNet50 CNN. 4,000 image tiles from each dataset were selected randomly to illustrate. Each point represents each feature vector of one image tile.

**Figure S5.**
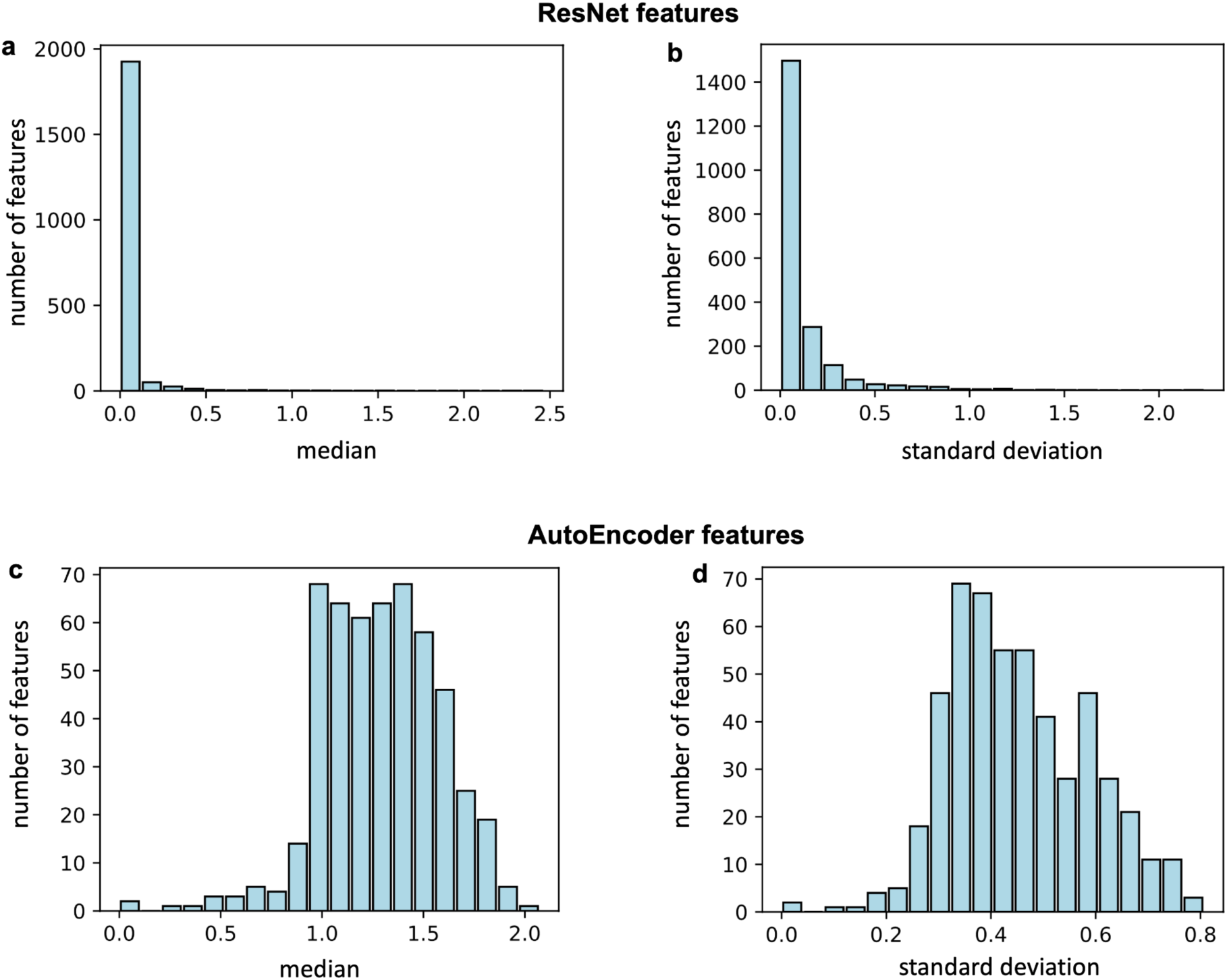
Histograms of median and standard deviation of ResNet features (upper) and AutoEncorder features (lower); Median (left panels) and standard deviation values (right panels) are shown. The TCGA-BRCA cohort was selected as an example.

**Figure S6.**
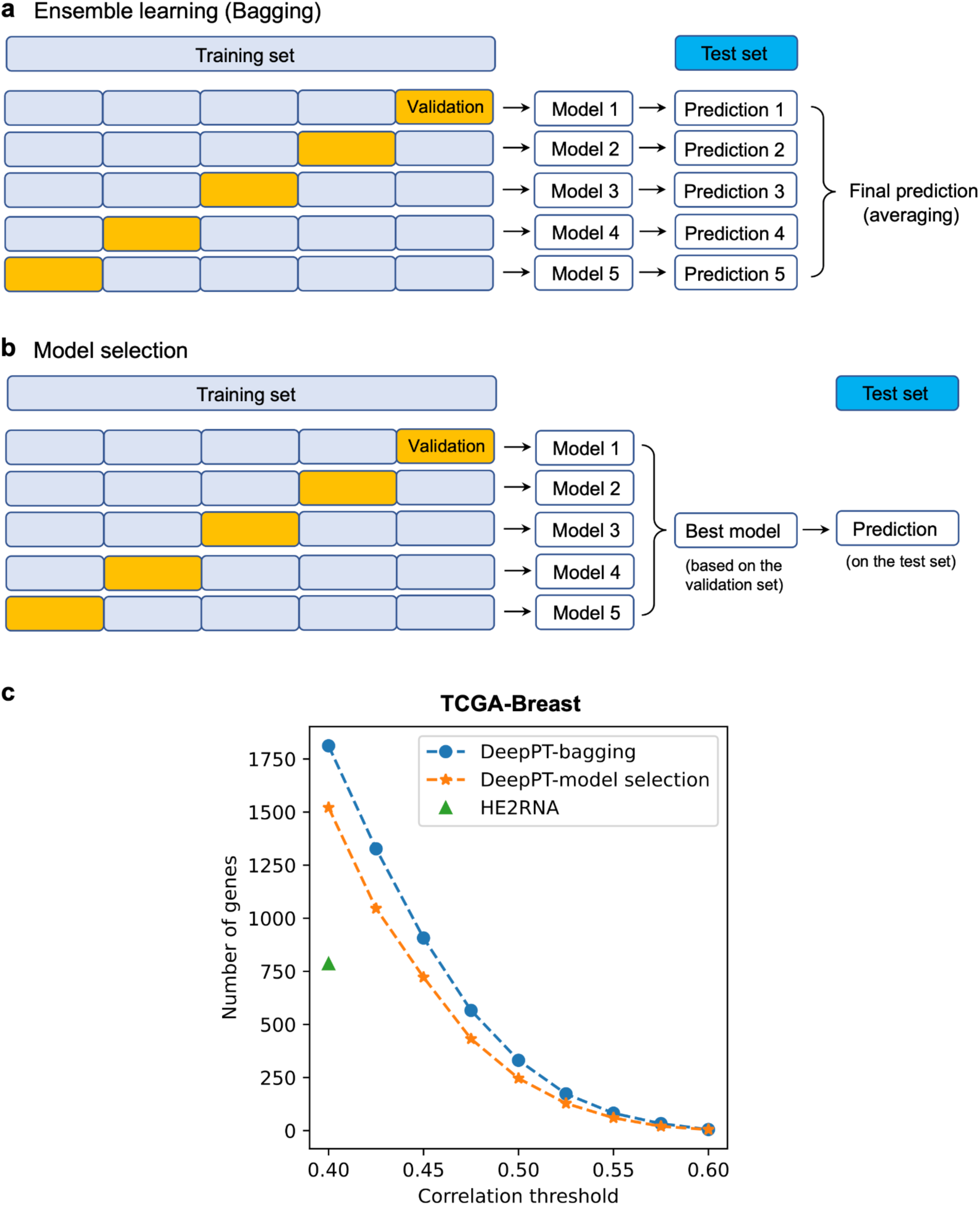
Training strategies and their performance. **(a)** In the ensemble learning strategy (bagging), five models were trained independently with five internal training-validation splits; these five model predictions were averaged to make the final prediction. **(b)** In the model selection strategy, the “best” model with the highest performance on the validation set was chosen to make prediction on the test set. **(c)** Number of genes with mean correlation over 5 folds greater than a certain threshold, obtained from these strategies. The TCGA-Breast cohort was selected as an example. Note that with either strategy, DeepPT outperforms the current state-of-the-art approach, HE2RNA.

**Figure S7.**
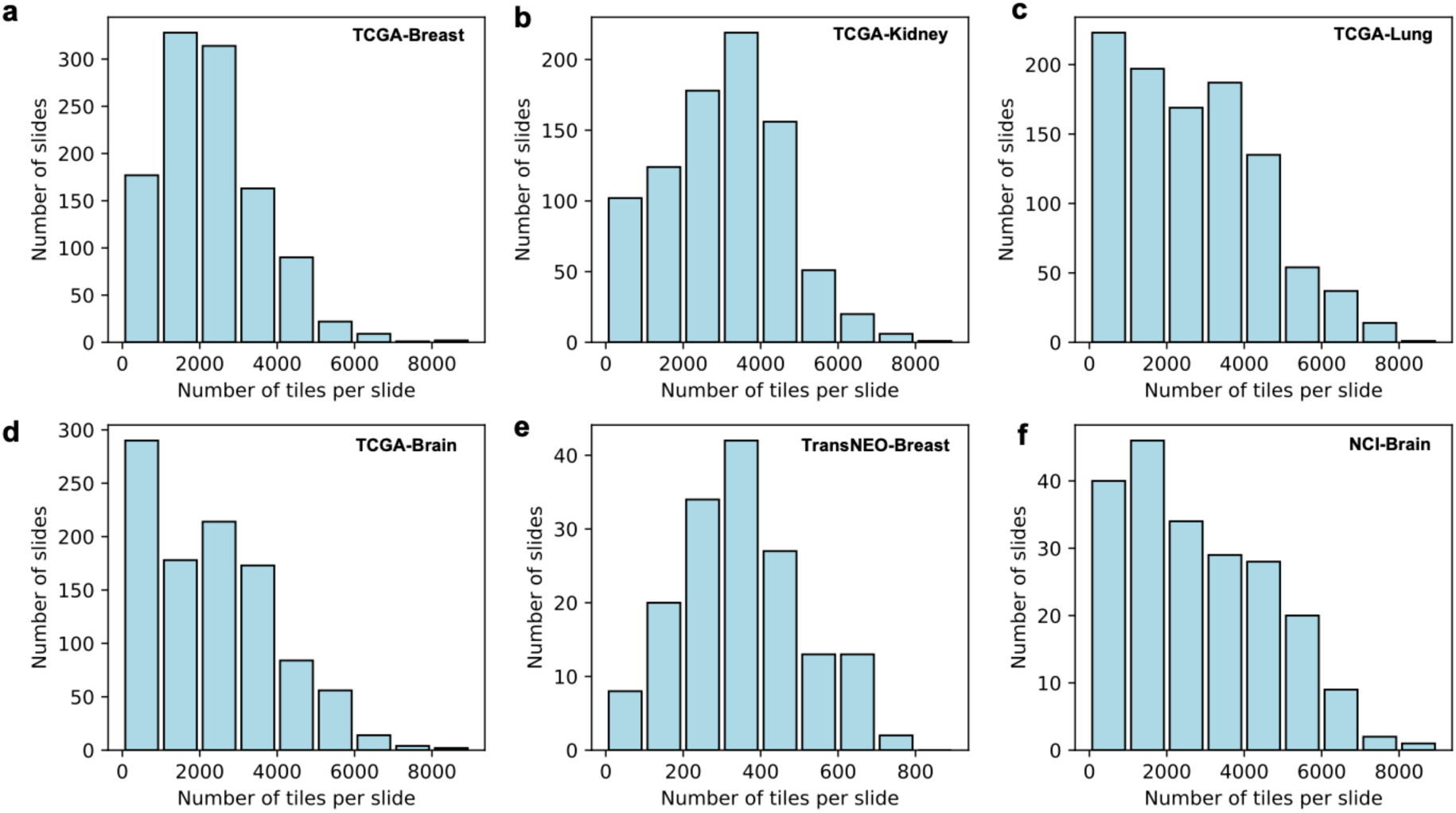
Histograms of the number of tiles per slide by cohort. The number of tiles in each slide image from TCGA and NCI-Brain datasets ranges from 100 to 8,000 (a, b, c, d), while the number of tiles in each TransNEO-Breast slide image is much smaller, ranging from 100 to 1,000 (e).

**Figure S8.**
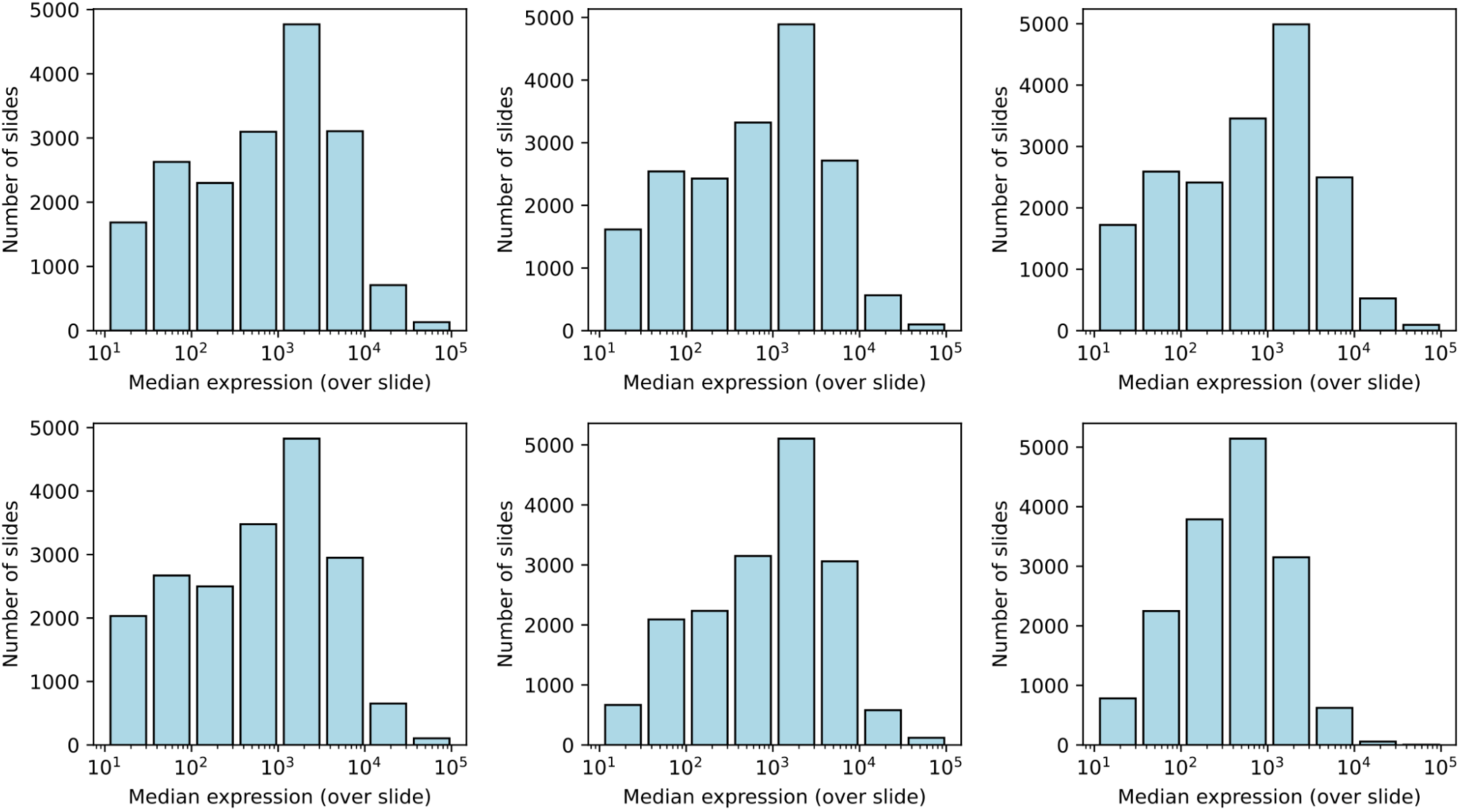
Histogram of median expression over slides. The median expression over samples of each gene commonly varies from 10 to 100,000 for every dataset considered in this study.

**Figure S9.**
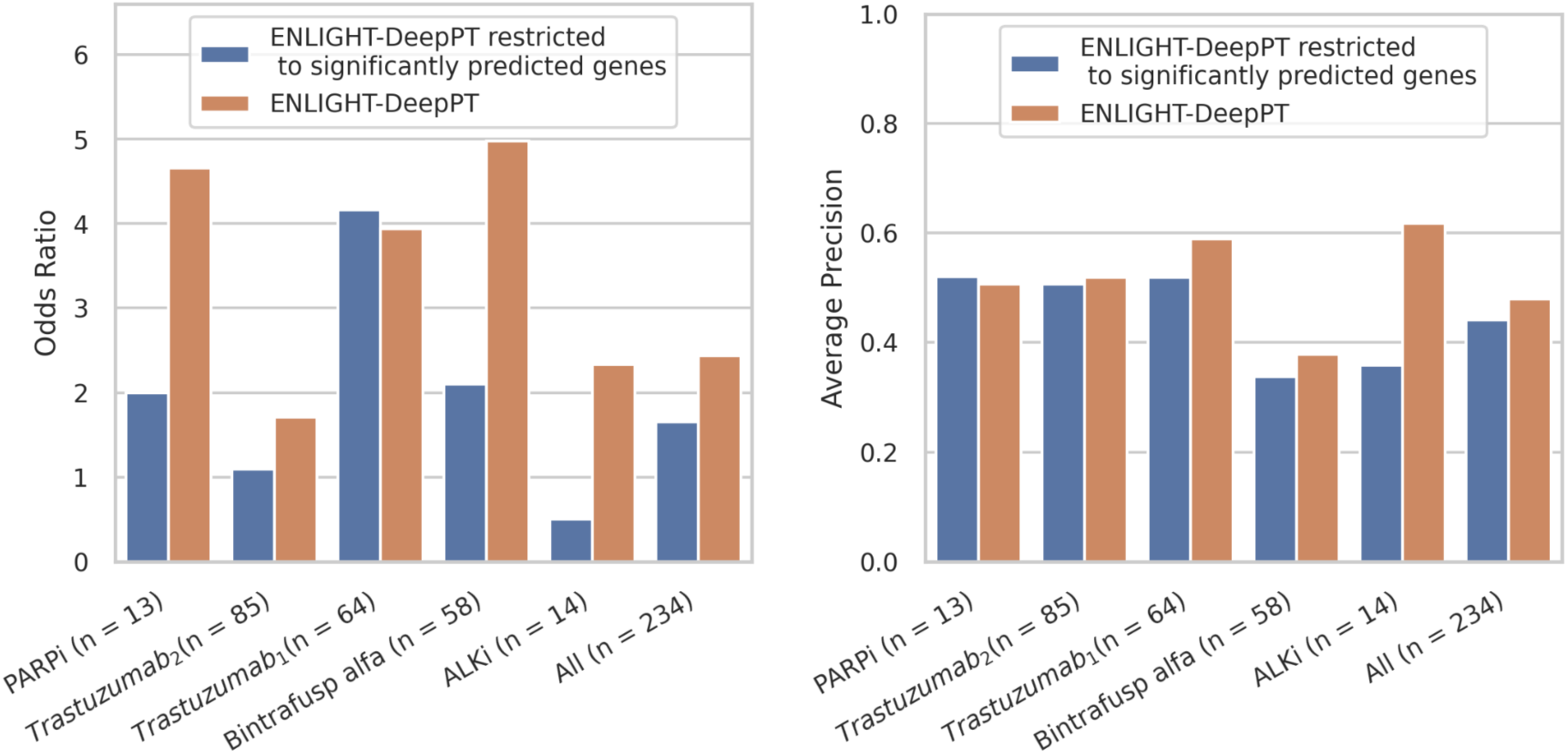
A comparison between the performance of ENLIGHT-DeepPT when using the same methodology described in [1] to generate genetic interaction networks that constitutes ENLIGHT’s predictive biomarkers (orange bars) and a revised methodology (blue bars) where we restricted ENLIGHT’s biomarker to only include genes that showed significant positive correlation (corrected p < 0.05) between actual and DeepPT-predicted values among the respective TCGA cohort (that is, according to the cancer type of each of the five drug response datasets). Left panel: Odds Ratio (OR) for each dataset, using the same clinical decision threshold that has been previously established in [1]. Right panel: Average Precision (AP) for each dataset.

**Figure S10.**
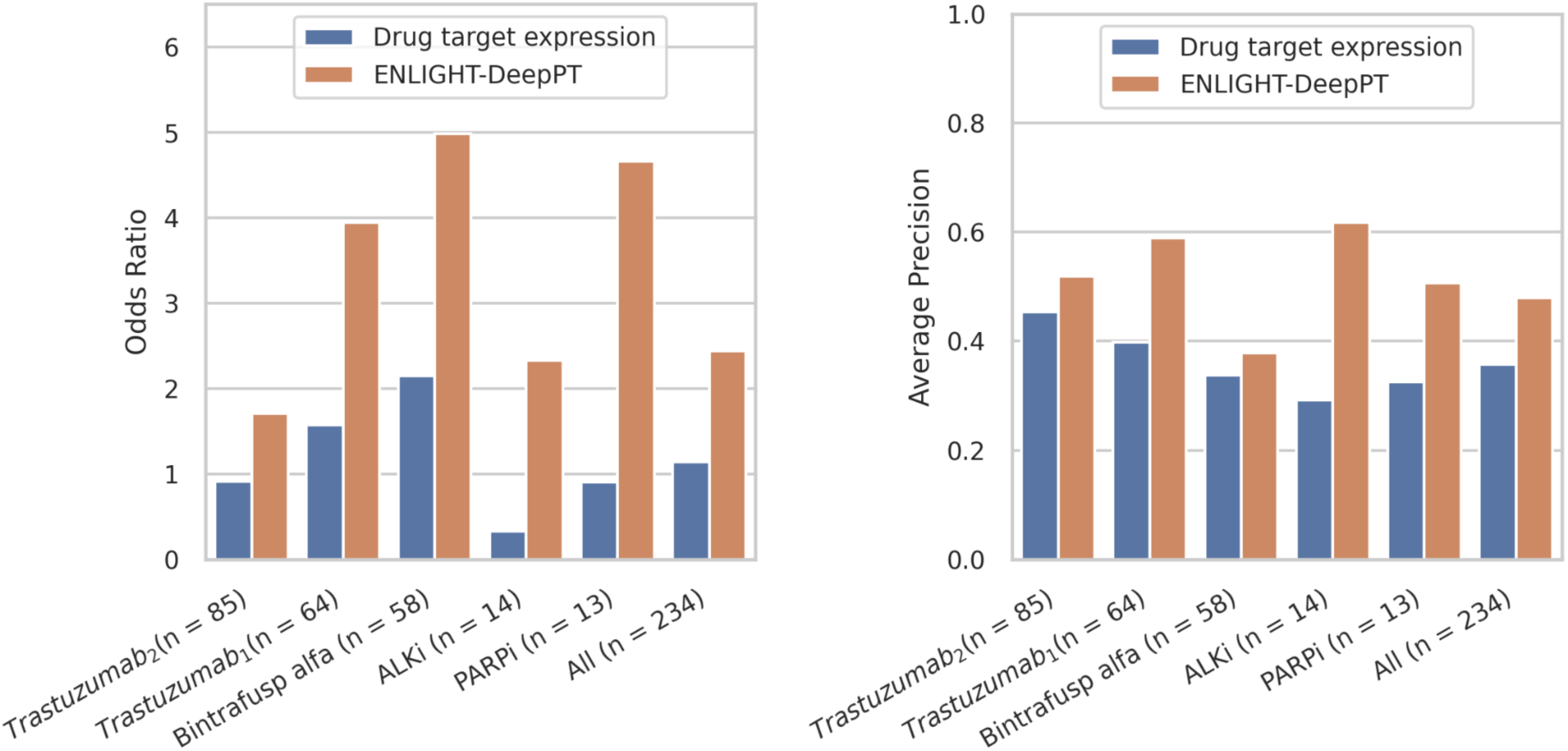
Comparison of the predictive performance of ENLIGHT-DeepPT (orange bars) and the respective drug target(s) expression models (blue bars) for each patient cohort and on the aggregation of all patients. Left panel: Odd ratio (OR) for each cohort for the same thresholds established in [1]: Average Precision (AP) for each cohort.

**Figure S11.**
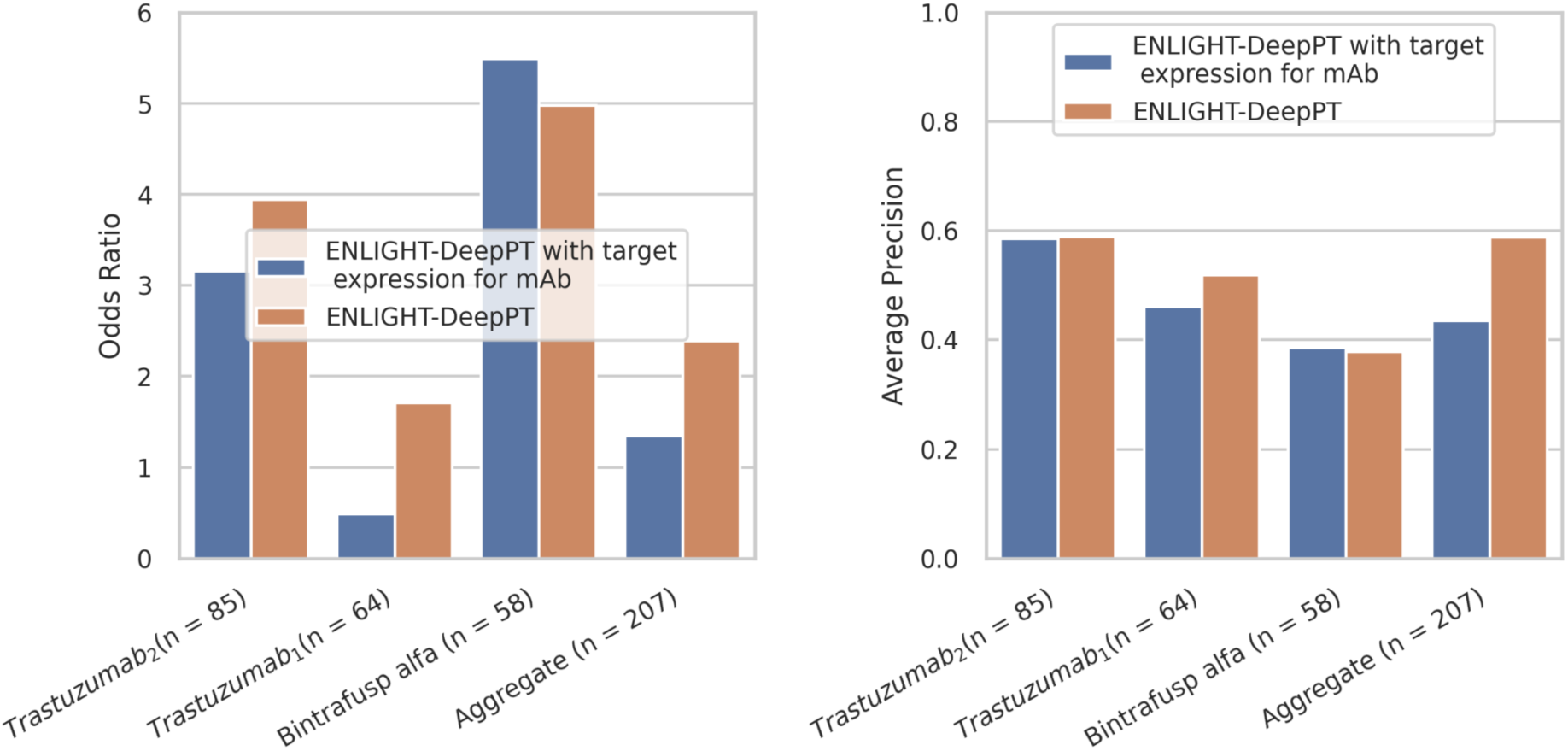
Comparison of the predictive performance of ENLIGHT-DeepPT (orange bars) and a version of ENLIGHT-DeepPT that incorporates the target expression in the scoring method for antibodies. Results are shown for each of the three dataset where antibody drugs were used and the aggregation of them. Left panel: Odd ratio (OR) for each cohort for the same thresholds established in [1]. Right panel: Average Precision (AP) for each cohort.

**Table S1.** Number of slides and number of patients for each cohort.

| **Cohort** | **#slides in total** | **#slides excluded** | **#slides remaining** | **#patient remaining** |
| --- | --- | --- | --- | --- |
| TCGA-Breast | 1,125 | 19 | 1,106 | 1,043 |
| TCGA-Lung | 1,042 | 24 | 1,018 | 927 |
| TCGA-Brain | 1,039 | 24 | 1,015 | 574 |
| TCGA-Kidney | 876 | 17 | 859 | 836 |
| TCGA-Colorectal | 614 | 100 | 514 | 510 |
| TCGA-Prostate | 445 | 7 | 438 | 392 |
| TCGA-Gastric | 508 | 75 | 433 | 410 |
| TCGA-Head&Neck | 444 | 14 | 430 | 409 |
| TCGA-Cervical | 277 | 16 | 261 | 252 |
| TCGA-Pancreas | 195 | 0 | 195 | 175 |
| TransNEO-Breast | 160 | 0 | 160 | 160 |
| NCI-Brain | 226 | 0 | 226 | 226 |

**Table S2.** Details of the 5 datasets analyzed using ENLIGHT-DeepPT. OS - overall survival; pCR - pathological complete response; RD - residual disease; CR – complete response; PR – partial response; RCB – residual cancer burden.

| Name | Indication | Drug | Background Therapy | N | # of responders | Responders criteria |
| --- | --- | --- | --- | --- | --- | --- |
| PARPi | Pancreas | Olaparib or Veliparib |  | 13 | 5 | OS > 36 months |
| Trastuzumab<sub>1</sub> | Breast | Trastuzumab | Chemotherapy | 64 | 19 | pCR |
| Trastuzumab<sub>2</sub> | Breast | Trastuzumab | Chemotherapy | 85 | 36 | pCR |
| Bintrafusp alfa | Lung, Head&Neck, Cervical | Bintrafusb alfa |  | 58 | 13 | CR, PR |
| ALKi | Lung | Alectinib or Crizotinib |  | 14 |  | > 18 months PFS |

